# Mechanotendography: description and evaluation of a new method for investigating the physiological mechanical oscillations of tendons using a piezo-based measurement system

**DOI:** 10.1101/2020.06.19.161174

**Authors:** Laura V Schaefer, Frank N Bittmann

## Abstract

The mechanotendography (MTG) analyzes mechanical oscillations of tendons during muscular actions. It can be assessed as equivalent to mechanomyography just applied for tendons. Since this method is unknown, the aim of this investigation was to evaluate the technical reliability of a piezo-based measurement system used for MTG.

The reliability measurements were performed using audio files played by a subwoofer. The thereby generated mechanical pressure waves were recorded by a piezoelectric sensor based measurement system. The piezo sensor was fixed onto the subwoofer’s coverage. An audio of 40 Hz-sine oscillations and, to stay close to human applications, four different formerly in vivo recorded MTG-signals from Achilles and triceps brachii tendon were converted into audio files and were used as test signals. Five trials with each audio were performed. One audio was used for repetition trials on another day. The correlation of the recorded signals were estimated by the Spearman correlation coefficient (MCC), the intraclass-correlation-coefficient (ICC(3,1)), Cronbach’s alpha (CA) and by mean distances (MD) between the signals. They were compared between repetition and random matched signals.

The repetition trials show high correlations (MCC: 0.86 ± 0.13, ICC: 0.89 ± 0.12, CA: 0.98 ± 0.03), low MD (0.03 ± 0.03V) and differ significantly from the random matched signals (MCC: 0.15 ± 0.10, ICC: 0.17 ± 0.09, CA: 0.37 ± 0.16, MD: 0.19 ± 0.01V) (*p* = 0.001 – 0.043).

This speaks for an excellent reliability of the piezo-based measurement system in a technical setting. Since research showed that the skin above superficial tendons oscillates adequately, we estimate this tool as valid for the application in musculoskeletal systems. It might provide further insight into the functional behavior of tendons during muscular activity.

## Introduction

Neuromuscular oscillations are frequently considered in medical, sports and health sciences. Thereby, the motor output is mostly examined by electromyography (EMG). Meanwhile, the method of mechanomyography (MMG) is also utilized more often. It is common knowledge that muscle fibers are mechanically oscillating in a stochastic way distributed in low frequency ranges of around 10 Hz [1-3]. The muscles function as actuators of the musculoskeletal system, which are mostly linked to the bones by tendons. Tendons are passive connective tissue structures, which are driven by the active parts – the muscles. However, the force during movement is transmitted from muscle via aponeurosis and tendon to the skeleton [4]. Thereby tendons have to transmit great forces. For instance Achilles tendons are strained with a tensile stress of about 9000 N during running [5-7]. Basic research concerning tendons focuses on the investigation of cadaver tendons, e.g., to get information about the distinct structures of tendons with different physiological requirements with regard to their function [8,9] or, respectively, on modelling as well as linear and non-linear analysis of the mechanics of tendons [10]. Clinical examinations in humans are usually performed by ultrasound to clarify the anatomical structure or changes of them [11,12]. Real-time ultrasound allows the scanning of the tendon during isometric contraction [13-17]. Thereby, the focus of interest lies on the force-displacement [15] and the deformation and stiffness [14] of the tendon during isometric contraction of the plantar flexors and thereby get information about stress and strain of the tendon [13]. An oscillatory displacement of the tendon is not considered thereby. The oscillations of tendons, which are generated during the muscular force transmission, are referred to as tendinous output, which is rarely investigated in research. The real-time ultrasound is used in lower sampling rates of 25 Hz [14], which is too low to analyze the tendinous oscillations. Established methods as MMG and EMG, respectively, can only provide information about the muscles, not about the passive structures. However, these passive structures often develop complaints as, e.g., tendinopathies. Since a change of motor control is inter alia discussed as potential factor for the development of tendinopathies [7,13,18-21], it would be of special interest to investigate the mechanical oscillating behavior of tendons, which are generated by the connected muscles. The recording of tendinous oscillations might provide further insights than only capture the muscular activity, since tendons often reflect the oscillations of more than muscle, e.g. the Achilles tendon, where three muscle heads insert. It also might reveal more information than looking at the mechanical tendinous structure as fibre structure and thickness, stiffness, displacement and strain during isometric contraction, since it includes the oscillatory behavior, which is generated by the neuromuscular system. Therefore, the recording of tendinous oscillations during motor actions might provide a more functional insight into the properties of tendons.

Recently, the tendon shear wave were assessed using high frame rate ultrasound, accelerometers or laser Doppler vibrometry after tapping on or shaking the tendon, respectively [22-25]. The main finding of Salman [24] was that with increasing contraction, the tendon stiffness increased.

The research group around Martin et al. [22**Error! Bookmark not defined**.] investigated the spread wave of the Achilles tendon using two accelerometers after tapping on the tendon during cyclic isometric ankle plantar flexion and during walking. Due to the change of shear wave in different gait phases, they draw the conclusion about the passive stretch of the tendon, e.g. before heel strike [22]. This is close to the investigation of [25], in which a myotonometer is used to apply a mechanical impact on the tendon and to record the resulting oscillation of the tissue in relaxed tendons. This technique is quite similar to the Supersonic shear imaging (SSI), which also generates vibrations of tissue (using ultrasonic beams) and examines the shear wave of the tissue afterwards [26]. In all those investigations, firstly, vibrations of the tendon are induced from external to measure the resulting shear waves of the tendon with different techniques. However, when active, the muscles tighten the tendon. In this process, the muscles produce stochastically distributed mechanical oscillations, which can be measured via dynamometry and kinematics also without applying an external vibration or tapping [3,27,28]. The oscillations work in the axial direction along the tendinous strand. Thereby, the muscles act as actuators stimulating the passive tendon longitudinally and laterally. As far as we have assessed, no investigation recorded those oscillations of the tendon during muscle activation. Only a combination of muscle action and application of mechanical vibrations or tapping to the tendon are regarded, whereby especially the evoked shear wave of the tendon was of interest so far. This focuses especially on gathering information about the mechanical behavior after tapping or the like, which changes depending on the structures. Due to the connection of neuromuscular dysbalances and tendon pathologies, a technique, which considers the oscillations, might reveal a more functional insight into the tendinous behavior during muscle activation. The oscillations of tendons during isometric muscle action can be detected by piezoelectric sensor based measurement systems, which will be presented here. We estimate this technique as unique and innovative. It is suggested to name this method mechanotendography (MTG). It can be considered as analogy to MMG but applied for tendons [3,28,29].

Because of the novelty of this method using a piezo-based measurement system, there is the need of evaluation. The measurement system is adopted from music. The piezoelectric sensors and amplifiers are usually used to pickup and amplify auditory signal from instruments. Thereby, they have been proofed to be suitable to take off harmonic oscillations. Anyway, the justifiable question remains, whether or not piezoelectric sensors are suitable to record stochastically distributed mechanical oscillations in frequency areas around 20 Hz like those produced by muscles or tendons.

Accordingly, the first step is to investigate whether or not this measurement system is able to capture low frequency oscillations reliably in a pure technical setting (ex vivo). This was the objective of this study. The basic applicability of MTG in vivo was already shown in several publications. MTG and MMG of the triceps brachii muscle and tendon are, e.g., able to generate coherent behavior during isometric muscle action of the elbow extensors [3,28]. Furthermore, MTG was applied in an investigation of patients with Achilles tendinopathy [30]. Detecting the tendinous oscillations using a low cost method like here presented might provide a more functional insight into the tendinous behavior during load.

## Materials and methods

Although in the presented study the measurements took place in a solely technical setting (ex vivo), the method will be called MTG in the following.

### Piezoelectric sensor and amplifier used for MTG

A large number of piezoelectric sensors were tested for the usage of recording the mechanical oscillations of tendons and muscles in the Potsdam Neuromechanics laboratory (Germany). Instrumental pickups are used to convert mechanical vibrations of solid objects into electrical voltage, which in the usual application results in an audio signal (tones of instruments). Mechanical pressure or structure-borne sound generate an electric voltage thereby, e.g. in wind instruments. Considering the tendon as string, the structure-borne sound generated by the tendinous swing can be recorded by piezoelectric pickups. However, not every pickup is applicable. Two pickups have been proven to be suitable to identify the mechanical oscillations of muscle-tendon structures: Shadow SH 4001, usually used as pickup for clarinets and saxophones, and the Shadow SH-SV1, a pickup for violins. For the present investigation, the piezoelectric sensor Shadow SH 4001 was used to verify the reliability of those sensors (in the following called MTG-sensor). One of the benefits of those sensors is the small size (ø12mm) (Figure 1).

**Fig 1.**
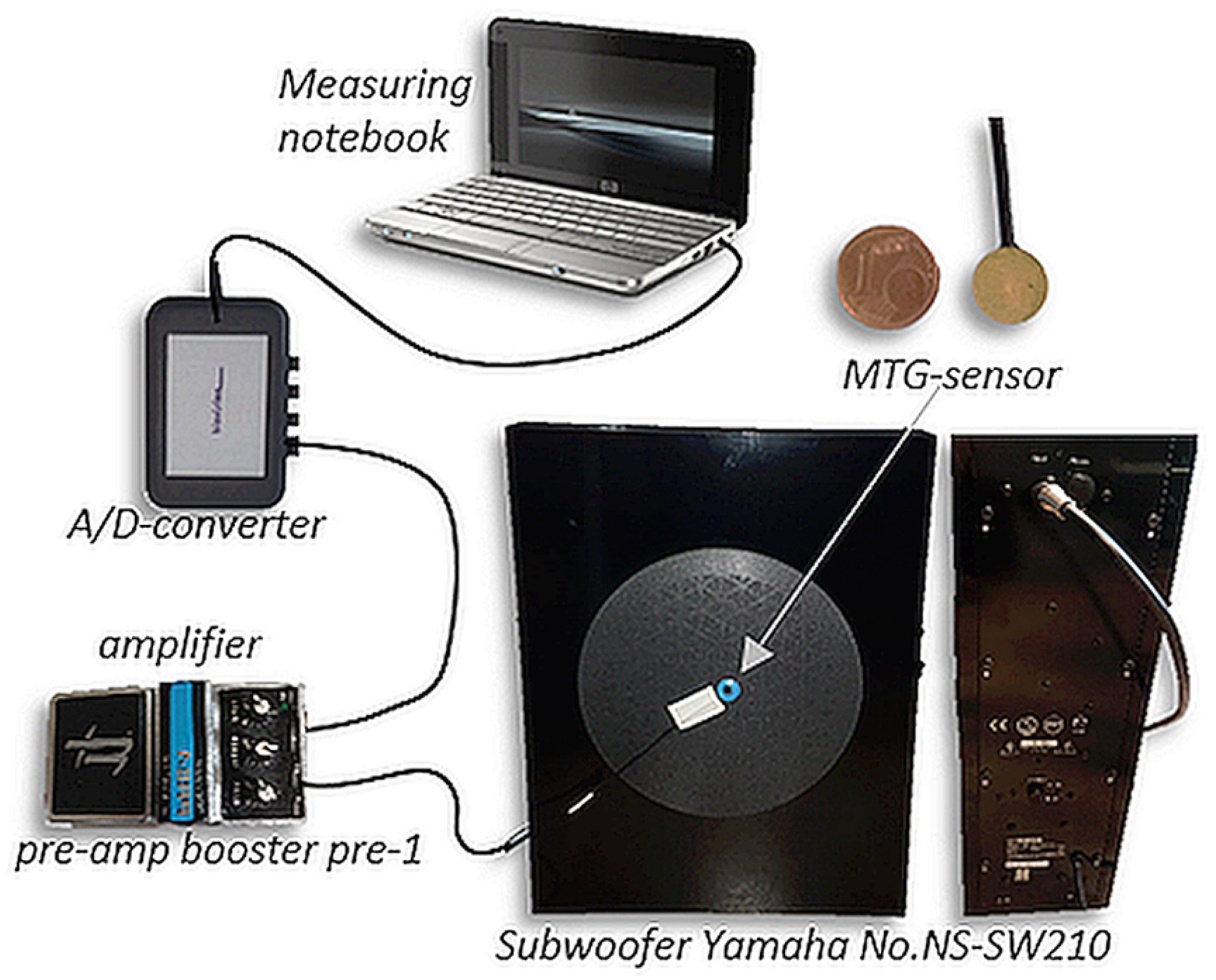
Setting. The piezoelectric sensor (Shadow SH 4001) was fixed on the coverage of the subwoofers’ loudspeaker (Subwoofer Yamaha No. NS-SW210) and was connected to an amplifier (Nobels pre-amp booster pre-1). The signal was transmitted through an A/D-converter to the measuring notebook (Lenovo V330). The signal was recorded with NI DIAdem 14.0.

However, the pickups are useless without an adequate amplifier. Therefore, the choice of an appropriate amplifier is essential. For MMG/MTG this has to be suitable to amplify low-frequency ranges below 30 Hz. The amplifier Nobels pre-amp booster pre-1, usually used for guitars, turned out to be applicable. Several other amplifiers were tested, but were not reproducing and amplifying the signal properly.

### Setting and procedure

In order to examine the reliability of the piezo-based measurement system, different audio sounds (origin see below) were played by the Software NI DIAdem 2017 (National Instruments) via a subwoofer. The subwoofer Yamaha No. NS-SW210 Advanced YST II was used to generate mechanical pressure waves, which were picked up by the MTG-sensor. The subwoofer has a frequency response from 30 to 160 Hz and, therefore, appears to be just appropriate to reflect low frequency ranges. The MTG-sensor was fixed by special ECG- and adhesive tape onto the coverage of the loudspeaker of the subwoofer, where the pressure waves should be the highest (Figure 1). This same fixation is also used for measurements in vivo. The picked up signal was amplified by the amplifier Nobels preamp booster pre-1 and was transferred via A/D-converter (NI USB-6218, 16-bit) to the measuring notebook, where it was recorded by NI DIAdem 14.0.

Five different audios were played by the software NI DIAdem via the subwoofer and were picked up. Thereof, three audios were converted from different original MTG-signals recorded during measurements of the Achilles tendon after impact (recorded in previous investigations on humans [30]): audioMTGAchilles_1, audioMTGAchilles_2, audioMTGAchilles_3. Furthermore, an original MTG-signal recorded from the triceps brachii tendon during isometric muscle action of the triceps brachii at an intensity of 80% of the MVC was converted into an audio file (audioMTGtri). They were used to produce a signal which is close to real application. The setting of capturing those signals in vivo can be looked up in [**Error! Bookmark not defined**.] and [**Error! Bookmark not defined**.], respectively. Additionally, a 40 Hz sine oscillation was produced by an online tone generator (audioMTG40Hz) to get further information concerning the amplitude and frequency reproducibility of the MTG-system.

To examine trial-to-trial reliability, five repetitions of each audio were played and recorded by the piezo-based measurement system at the same day. The audioMTGAchilles_1 was re-recorded one day later (5 trials) to investigate the day-to-day reliability. To investigate the objectivity and validity, the audio of 40 Hz sine oscillations was used.

### Data processing and statistical analysis

One part of the data consideration was the visual descriptive evaluation of repeated trials. For further data evaluation, the raw data of each curve were used. Each recorded MTG-signal was cut into a short interval of 0.1s (MTGAchilles) and 0.5s (40Hz-sine, MMGtri), respectively using NI DIAdem 2017. The 0.1s interval of MTGAchilles-audio was chosen because the original MTG-signal (in vivo) was recorded during a short impact on the forefoot of the participant from plantar in direction of dorsiflexion, which generated the here relevant oscillation of the Achilles tendon in the 0.1s interval after impact [30]. The original MTGtri-signal was originally recorded in vivo during a 10s isometric muscular interaction of two participants at 80% of the MVC (similar to arm wrestling) [3]. Since the sampling rate was 1000 Hz, 0.5s provides a sufficient long signal for the investigation of reliability. The measurement parameter in the present study (ex vivo) was the amplified voltage of the piezoelectric sensor gathered from the mechanical oscillations of the subwoofer during audio replay. To investigate the reproducibility, the repetition trials of each audio were compared between the trials by analyzing the following parameters: (1) mean distances of all data points between the curves (MD) in Excel 2016 and (2) the spearman correlation coefficient (MCC), (3) the Intraclass correlation coefficient (ICC(3,1)) and (4) Cronbachs alpha (CA) in IBM SPSS 25. Concerning the MD (1) and MCC (2), ten values arise from the five trials for each signal. For further statistical comparisons, therefore, the arithmetic mean (M) and standard deviation (SD) were calculated.

To compare these items to randomized matched curves, MTG-signals of the three different MTGAchilles-audios were used to form five random groups. Only signals of the MTGAchilles audios were chosen, since the 40 Hz-sine and the MTGtri-audio show very different characterization compared to the MTGAchilles-signals and would skew the results significantly. In human investigations, the MTG is used to compare similar settings, therefore, a MTG of the Achilles tendon after impact, a MTG of the triceps brachii muscle during isometric muscle activation and 40Hz-sine oscillations would not be compared directly. Therefore, the randomized groups only contained trials of the MTG-audios of the same setting. The above listed parameters (1) to (4) for estimation of the reliability were also calculated for the randomized groups.

Using IBM SPSS 25, the parameters (1) to (4) were tested concerning normal distribution using the Shapiro-Wilk-test in each group. In case of normal distribution, the t-test for dependent samples was used to compare the identical repetition groups to the random ones statistically (MCC, ICC). The data of the other parameters (MD, CA) were not normally distributed. Therefore, the non-parametric Wilcoxon-test was used for those comparisons. The tests for dependent samples were chosen, since the random group also contains trials of the identical audio groups, and hence, the groups are not completely independent.

## Results

To illustrate the reproducibility of the oscillation characteristics of the original MTG-signals and the recorded corresponding audio-signal from this investigation, Figure 2 shows exemplarily one original MTGAchilles-signal and the corresponding recorded signal in the present setting using the subwoofer with fixed MTG-sensor. As shown, the frequency is reproduced precisely, the amplitudes differ.

**Fig 2.**
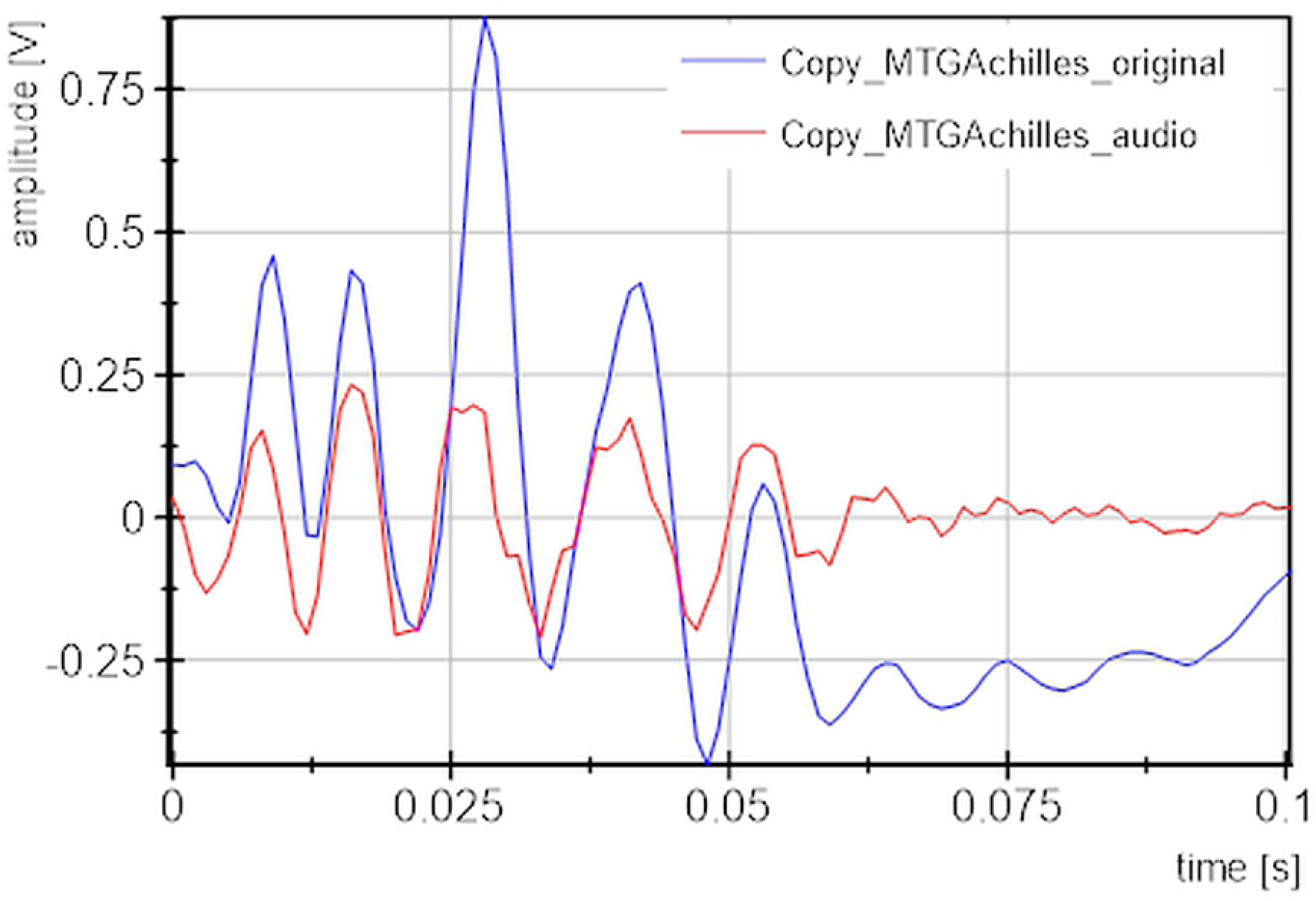
Exemplarily signals of the original MTGAchilles_1 signal (blue) compared to the MTGAchilles_audio (red) played by the subwoofer and recorded by the piezo-based measurement system in the present technical setting.

The curve shapes of the identical recorded trials are shown in Figure 3. In each diagram, five or, respectively, ten repetitions are displayed and reveal a good reproducibility. The day-to-day trials of the audioMTGAchilles_1 are displayed in Figure 4. As can be seen, the 10 signals lie highly reproducible one above the other. For quantification of this, the parameters (1) to (4) are regarded.

**Fig 3.**
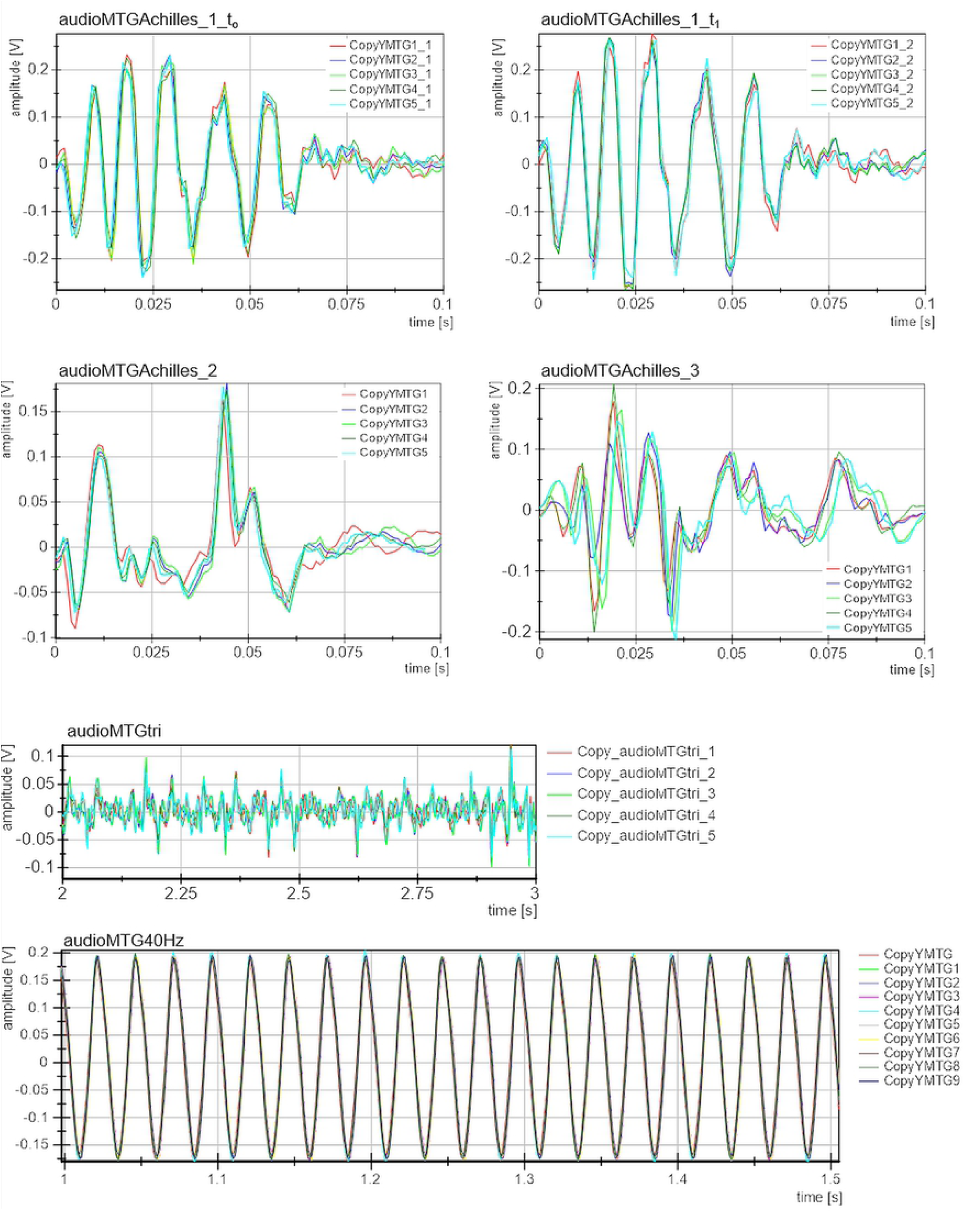
Displayed are the recorded MTG-signals by the piezo-based measurement system from the replay of the subwoofer. Each five trials of the same MTG audio and, respectively, 10 repetition trials of the 40Hz sine audio are illustrated in one diagram.

**Fig 4.**
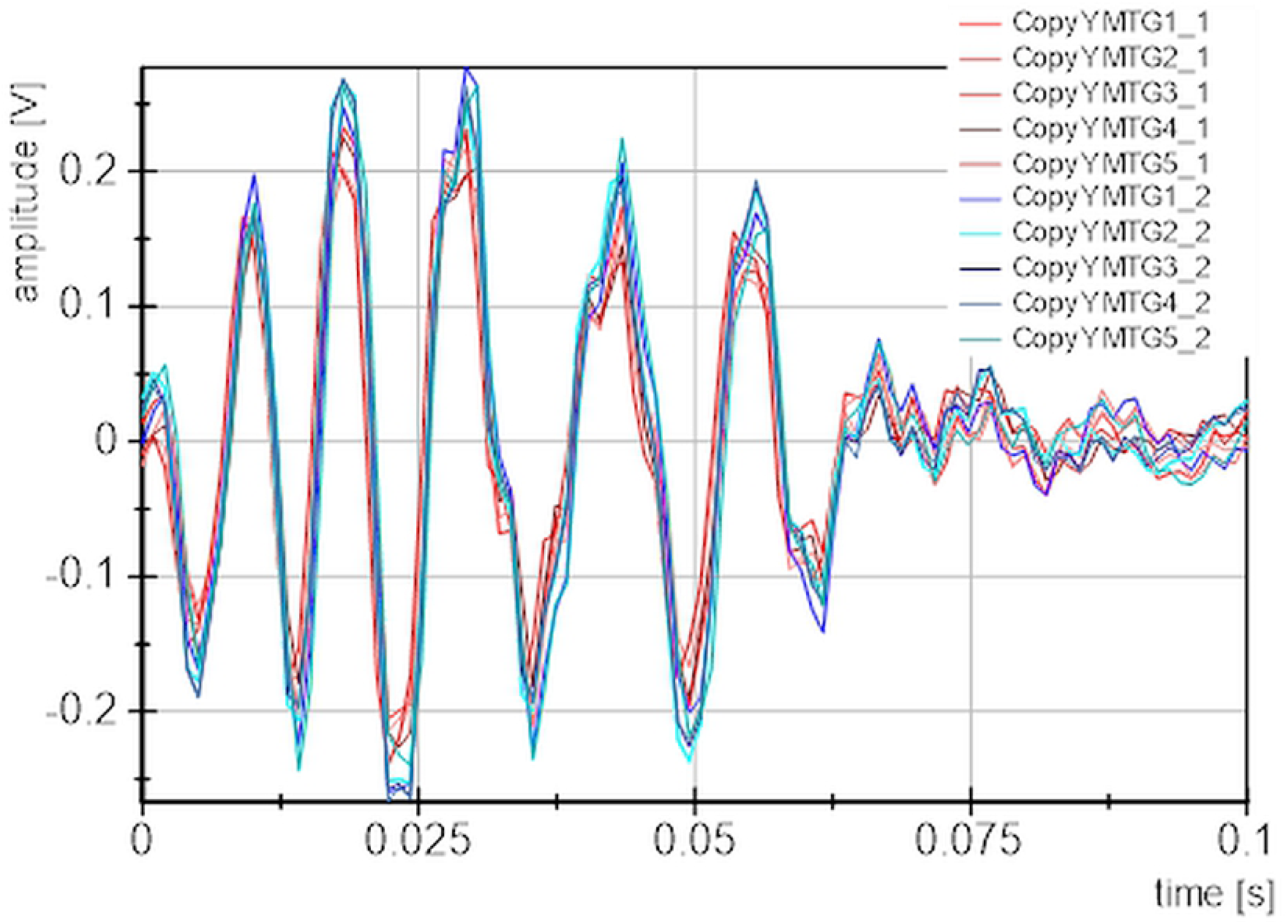
Displayed are each five repetition trials of the audioMTGAchilles_1 recorded with the piezo-based measurement system on two separate days (red colors = t_0_; blue colors = t_1_).

### Mean distances and correlation of repetition trials

The mean distances, mean spearman rank correlation, ICCs and Cronbachs alpha between the groups of the identical repetition trials and the random matched group are displayed for each recorded audioMTG-signal in Table 1. The comparisons of group averages (M ± SD) are illustrated in Figure 5.

**Table 1.**
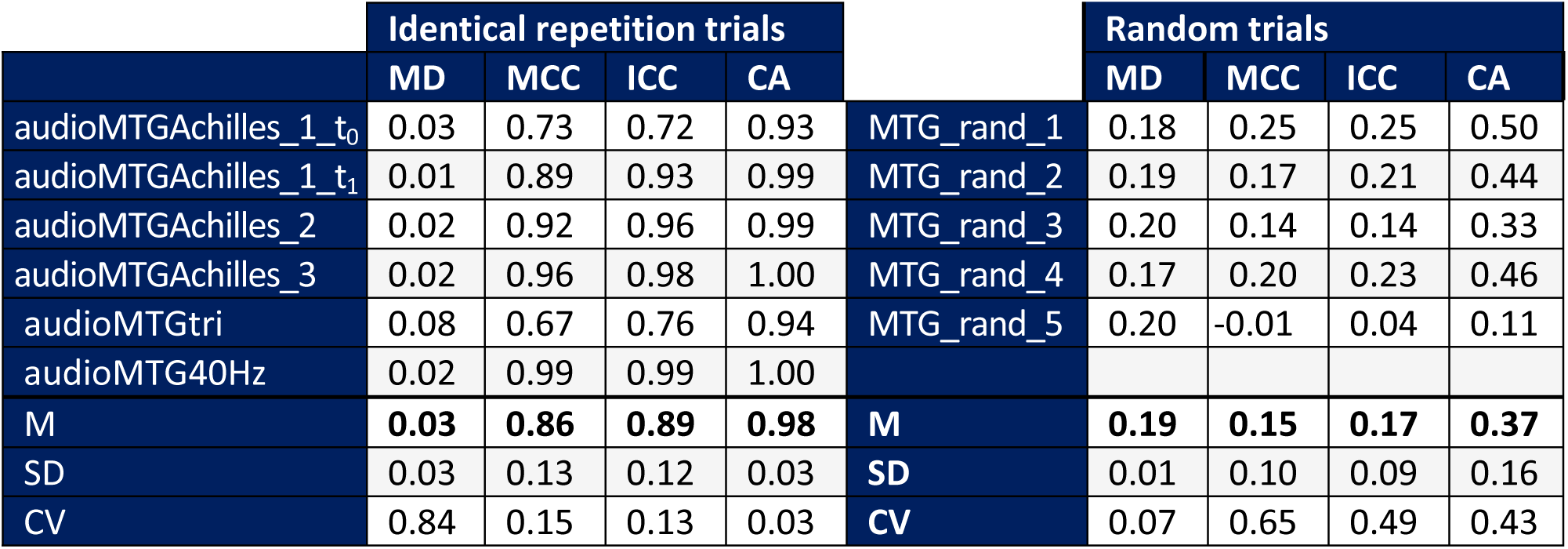
The parameters averaged mean distances (MD), mean spearman correlation coefficient (MCC), Intraclass correlation coefficient (ICC(3,1)) and Cronbachs alpha (CA) calculated between each 5 identical repetition trials for each MTG-audio and between random matched signals (MTG_rand) are displayed. Arithmetic mean (M), standard deviation (SD) and coefficient of variation (CV) are given.

|  | Identical repetition trials |  |  |  |  | Random trials |  |  |  |
| --- | --- | --- | --- | --- | --- | --- | --- | --- | --- |
|  | MD | MCC | ICC | CA |  | MD | MCC | ICC | CA |
| audioMTGAchilles_1_t<sub>0</sub> | 0.03 | 0.73 | 0.72 | 0.93 | MTG_rand_1 | 0.18 | 0.25 | 0.25 | 0.50 |
| audioMTGAchilles_1_t<sub>1</sub> | 0.01 | 0.89 | 0.93 | 0.99 | MTG_rand_2 | 0.19 | 0.17 | 0.21 | 0.44 |
| audioMTGAchilles_2 | 0.02 | 0.92 | 0.96 | 0.99 | MTG_rand_3 | 0.20 | 0.14 | 0.14 | 0.33 |
| audioMTGAchilles_3 | 0.02 | 0.96 | 0.98 | 1.00 | MTG_rand_4 | 0.17 | 0.20 | 0.23 | 0.46 |
| audioMTGtri | 0.08 | 0.67 | 0.76 | 0.94 | MTG_rand_5 | 0.20 | -0.01 | 0.04 | 0.11 |
| audioMTG40Hz | 0.02 | 0.99 | 0.99 | 1.00 |  |  |  |  |  |
| M | **0.03** | **0.86** | **0.89** | **0.98** | M | **0.19** | **0.15** | **0.17** | **0.37** |
| SD | 0.03 | 0.13 | 0.12 | 0.03 | SD | 0.01 | 0.10 | 0.09 | 0.16 |
| CV | 0.84 | 0.15 | 0.13 | 0.03 | CV | 0.07 | 0.65 | 0.49 | 0.43 |

**Fig 5.**
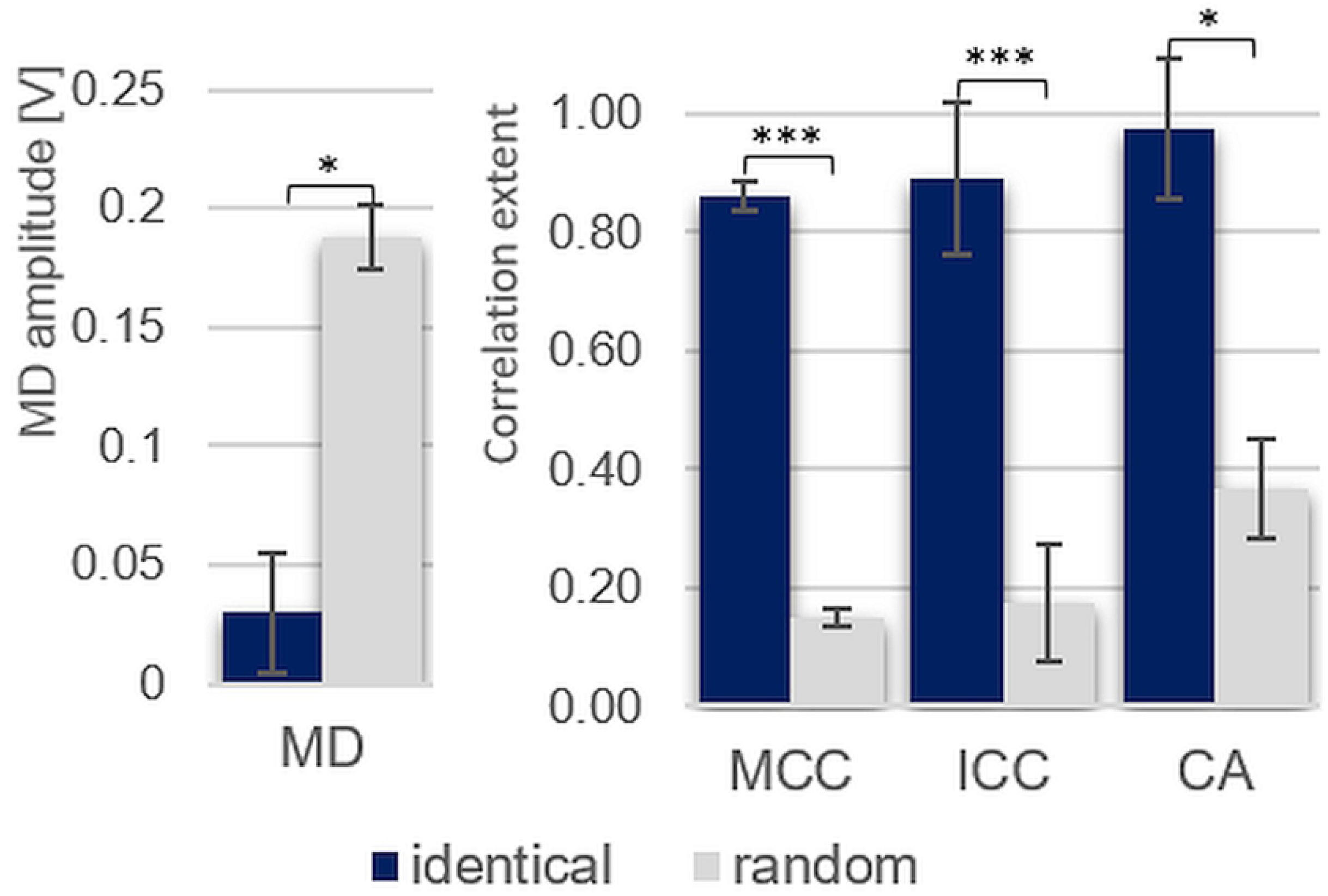
Arithmetic mean and standard deviation of the parameters mean distances (MD), mean spearman correlation coefficient (MCC), intraclass correlation coefficient (ICC(3,1)) and Cronbachs alpha (CA) of the identical repetition trials (blue) compared to the random matched group (orange). \**p* = 0.043, \*\*\**p* = 0.001

Looking at the correlation coefficients (MCC, ICC, CA) it is visible that the identical repetition trials have values from 0.67 to 1.0, which indicate good to excellent reliability [31], whereas the random matched groups show values from -0.01 to 0.25, which indicate no or poor reliability [**Error! Bookmark not defined**.] (Table 1). This is further supported by the group comparisons: The statistical comparisons between identical and random groups concerning the MCC and ICC show very high significance with t(4) = 9.104 (*p* = 0.001, *r* = 0.977) and t(4) = 9.317 (*p* = 0.001, *r* = 0.978), respectively. The Cronbach’s alpha show lower, but still significant differences (*W* = 2.023, *p* = 0.043, *r* = 0.905). These results indicate a significantly higher correlation of the identical repetition groups compared to the random matched ones.

In contrast, the parameter MD behave inversely: The averaged mean distances are significantly lower in the identical repetition group with mean values of 0.03 ± 0.03 V compared to the random matched group with an averaged MD of 0.19 ± 0.01 V (*W* = -2.023, *p* = 0.043, *r* = 0.905); reflecting the smaller distances between the curves of the identical repetition trials compared to the random matched signals. The distance between random matched curves are more than six times higher than the identical repetition trials.

For each comparison a high effect size of *r* > 0.90 is obtained, which underlines the significant differences between the identical and random groups.

## Discussion

The results show that the identical repetition group has significantly higher correlation values and considerably lower mean distances between the trials compared to the random matched group. Both results point out, that the curves during repetition of the same audio signal behave similarly. This indicates a high reliability of the piezo-based measurement system ex vivo.

The random matched curves behave inverse: The correlation parameters show, if at all, low correlation values and the mean distances are more than six times higher compared to the repetition curves of identical audio signals. This indicates a good distinction between different signals.

The discussion should focus on the technical aspects of the reliability measures and on the meaning of those for the application in vivo.

### Reproducibility of MTG-signals in the technical setting

The comparison of the original audio signals and the recorded signals with the piezoelectric sensor, which is exemplarily displayed in Figure 2, demonstrates a good agreement of wavelength, which indicates a high reproducibility of the frequency. The amplitude is lower in the recorded audio signal. Reasons for this lie probably in the replay of the subwoofer. The signal also shows partly signs of distortion. Because the undistorted original signals were detected in vivo by the same sensor type, the sensor can be excluded as the origin of this phenomenon. Therefore it is likely produced by the subwoofer, especially by the speaker’s coverage which is not as elastic as the speaker itself. Furthermore, especially in low frequency areas below 30 Hz, the subwoofer is not able to reproduce the sounds adequately. Since the mechanical oscillations of muscles and tendons during isometric muscle activity are to be found in those low ranges [1,2,3], the audios played by the subwoofer might reflect those frequency areas not as good as the original MTG-signals. Since the MTG-signals of the in vivo measurements of the Achilles tendon after impact show higher frequencies [30], they were chosen for the present reliability investigations, having in mind, that the subwoofer is able to reproduce them more precisely.

The limited frequency response of the subwoofer is especially visible in comparing the original MTG-signal of the triceps brachii tendon and the related recorded audio MTG-signal using the subwoofer (Figure 6). It is clearly visible that the signals do not match. This is led back to the low frequencies of about 15 Hz of the MTGtri-signal. The repeated recordings of this MTG-audio signal, however, indicate an only just excellent reliability *ICC(3,1)* = 0.76 (*p* = 0.000) [31]. It is concluded that the reproducibility of five repetition trials is very good, although the subwoofer is not able to rebuild the MTGtri-signal appropriately due to technical limitations of the frequency response of the subwoofer. It is assumed that with another subwoofer, which reproduces the low frequency ranges properly, the original and the recorded signals would be as similar as it was to be found for the MTG-signals of the Achilles tendon after impact. However, usually subwoofers are not required to play frequency ranges below 20 Hz, since they are not hearable for humans. Infrabass subwoofers would be able to reproduce such frequency ranges, but are only used in the professional event areas and are very expensive. Therefore, an infrabass subwoofer was not applicable in this setting. The aim of showing the reproducibility of repetition trials still is reached by the results of recording the MTGtri played by the subwoofer and recorded by the piezo-based measurement system from the coverage of the loudspeaker.

**Fig 6.**
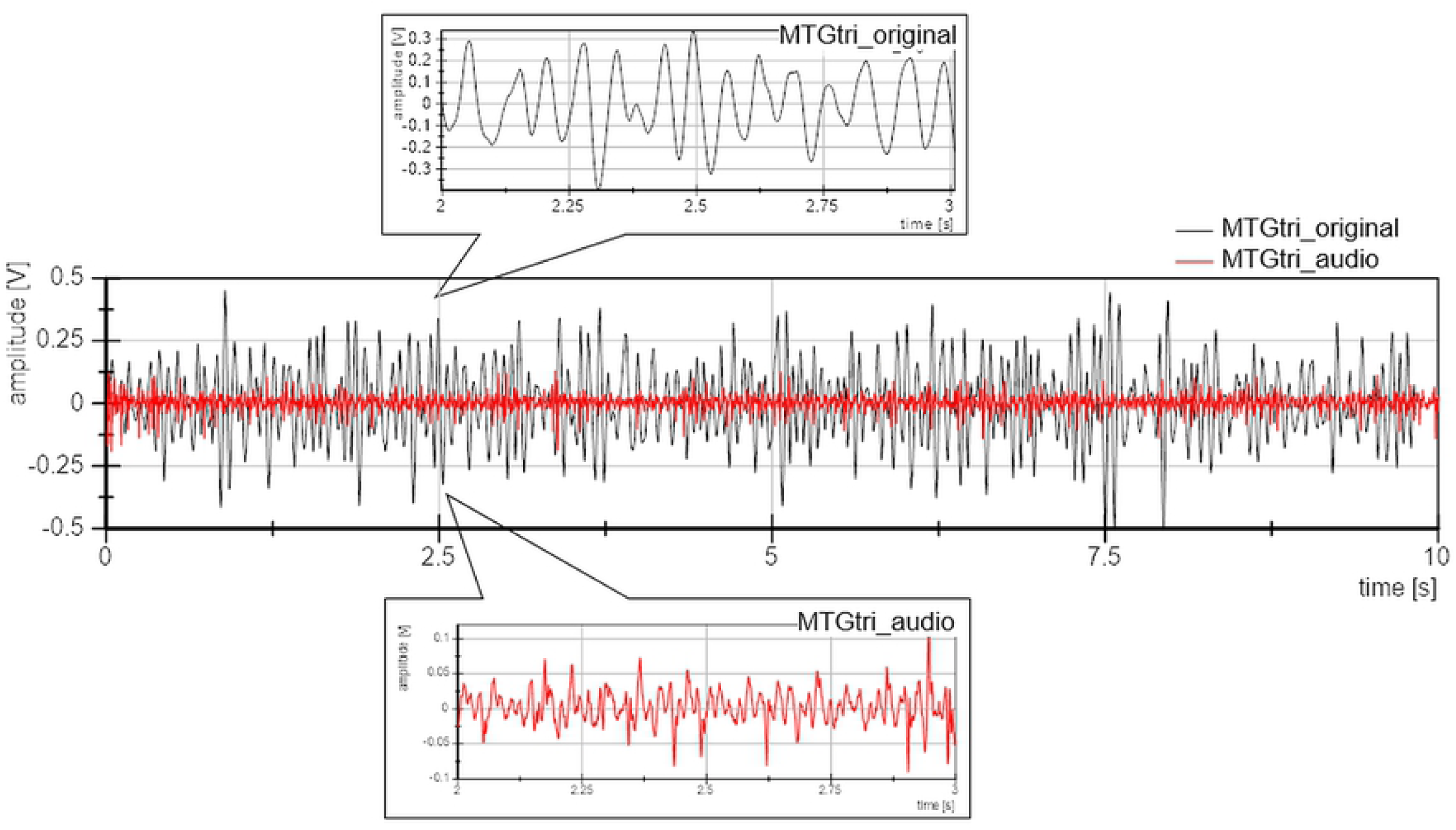
Comparison of the original MTG-signal of the triceps brachii tendon measured in vivo (black) to the corresponding recorded audio signal using the technical setting (ex vivo) (red).

### Reliability of repeated measurements using the piezo-based measurement system

The best reproducibility and, therefore, reliability was found for the technical 40 Hz oscillations (*ICC(3,1)* = 0.99, *p* = 0.000, *MCC* = 0.99), assuming that the piezoelectric sensor based measurement system is suitable to record harmonic oscillations properly in amplitude and frequency from the mechanical pressure waves of a subwoofer. The oscillations produced by a muscle or tendon is rather comparable to inharmonious structure-borne sound. As indicated here, the reproducibility of a former biological inharmonious signal is also captured in a reliable way using the subwoofer. Therefore, it can be concluded that the piezoelectric sensor based measurement system is a valuable tool to record mechanical pressure waves or structure-borne sound. This is not surprising with regard to the common application of those sensors in music. If the piezoelectric sensor would not reproduce the mechanical pressure waves in frequency and amplitude appropriately, the sound would be distorted. But what are the benefits comparing piezoelectric to acceleration sensors, which are mostly used for such investigations [1,22,32].

### Advantages of piezoelectric sensors used for MTG / MMG

From experiences in the Neuromechanics Lab, piezoelectric sensors, such as pickups, reflect the mechanical oscillations very precisely with an exceptional good signal to noise ratio (SNR). This is visible in Figure 6. The original signal of MTGtri was recorded from the triceps brachii tendon during isometric muscle activity of the elbow extensors with an intensity of 80% of the MVC. The raw data are displayed (Figure 6), thus, no filtering or smoothing were performed. The signal show clear, almost sinusoidal amplitudes in a frequency of around 15 Hz. None of the acceleration sensors we tested could reproduce such clear signals directly from the muscle belly or the tendon during isometric action.

However, not all piezoelectric sensors are suitable for the use of MMG or MTG. We presented here the – in our experience – most appropriate ones. However, we already had a new batch of the Shadow SH 4001 sensor, which have changed in quality. Therefore, we switched to the Shadow SH-SV1, which proofed to be suitable. There are other pickups, which turned out to be suitable with regard to the SNR. However, the one we tested had a larger diameter and, therefore, turned out to be not as practicable for fixing onto the skin above the muscle belly or tendon. Beside the choice of a suitable piezoelectric sensor, an essential factor is the used amplifier. As mentioned in the method setting, for MMG and MTG there is the need of an amplifier, which is capable to amplify low frequency ranges, too. Since the Nobels preamp booster pre-1 turned out to be suitable and reveals extremely clear signals, they are used in our investigations.

### Comparison of repeated audio trials to MTG trials in vivo

The piezoelectric sensor was already used in several studies in vivo for mechanotendography. In a study, the coherence of MMG and MTG-signals of the triceps brachii muscle and its tendon was investigated during muscular action of one person and muscular interaction between two persons (close to arm wrestling) during 80% of the MVIC [3]. Thereby it was shown that the MMG-/MTG-signal pairs of one measurement develop high coherence between the muscle and tendon of one person and also between two interacting persons. This was evaluated by wavelet coherence analysis. In contrast, random matched pairs did show significantly lower coherence [3]. In case the system would produce random, distorted or noisy signals, a result like this would not appear. This also indicates that the used piezo-based measurement system is a valuable and valid tool to measure tendinous and muscular oscillations in vivo.

The comparison of the here presented reliability results in the technical setting to measures of a recently conducted study in patients with Achillodynia [30] could lead to further assumptions. The MTG of the Achilles tendon was measured during an impact on the forefoot from plantar in direction of dorsiflexion during one leg stance (5 trials). Patients with Achillodynia (*n* = 10) showed a significantly higher mean spearman rank correlation (MCC) and a significantly lower averaged mean distance (MD) between the curves of five trials compared to healthy controls (*n* = 10) [30]. The MD amounted to 0.128 ± 0.029 V in patients and 0.227 ± 0.118 V in healthy controls (*p* = 0.028) and the MCC was 0.845 ± 0.073 compared to 0.451 ± 0.392 in healthy controls (*p* = 0.011). For the present reliability investigation, exemplarily signals from those investigations were converted into audio signals and were recorded. In comparing the results, it is even more indicated that the oscillations of an affected Achilles tendon after impact during repeated trials in vivo behave similarly compared to the repetition trials ex vivo recording the same audio file (MCC: Achillodynia (in vivo): 0.85 ± 0.07 vs. repeated audioMTGAchilles trials (ex vivo): 0.88 ± 0.13; MD: Achillodynia: 0.13 ± 0.03 vs. audioMTGAchilles: 0.02 ± 0.03). In contrast, the healthy subjects of the Achilles tendon study showed a higher variability indicated by a lower MCC and a higher MD. This behavior is rather comparable to the here presented random matched group (MCC: healthy Achilles tendon (in vivo): 0.45 ± 0.39 vs. random audioMTGAchilles (ex vivo): 0.15 ± 0.10; MD: healthy Achilles tendon: 0.23 ± 0.12 V vs. random audioMTGAchilles: 0.19 ± 0.01 V). This comparison between MTG-signals of the Achilles tendon in vivo and the recordings of the audio MTGAchilles-signals ex vivo indicate that immediately after an impact a healthy preloaded Achilles tendon oscillates in a more variable way, which tends to behave like random matched trials. Affected Achilles tendons (achillodynia) oscillate in a way, which rather matches the behavior of repeatedly recorded similar audio-signals of MTGAchilles, which show excellent reliability. It is assumed, therefore, that the higher variation in healthy controls is due to a necessary biological variability, which obviously plays an important role in healthy neuromuscular systems.

Because of their natural variability, biological systems will never produce identical wave patterns. Therefore, the validation of the reliability of such systems in vivo is limited. However, the high reproducibility of the tendinous oscillations in patients with achillodynia indicates that the piezo-based measurement system is not only suitable to capture the oscillations of audios in the here presented technical setting in a reproducible way, but also are able to monitor the oscillations of muscle and tendon oscillations reliably.

### Tendinous oscillations as possible insight into motor control

It is suggested that the mechanical oscillations captured superficially by the piezoelectric sensors reflects the motion of tendons. This is supported by the investigations of [22], in which it was shown by real-time ultrasound that the motion of the Achilles tendon and the adjacent subcutaneous tissue were similar. Therefore, it is conceivable that the mechanical pressure waves, generated by those oscillations, can be captured by the piezoelectric sensors fixed on the skin. Tendons oscillate laterally and axially. The piezoelectric sensor is not able to display the axial motion due to the placement on the skin. However, using MTG, the mechanical pressure waves, which are produced by a three-dimensional motion, can be captured in the transversal plane.

A special feature regarding some tendons, e.g. the Achilles tendon or the tendon of the triceps brachii muscle, is that there is more than one head, which inserts into the tendon. Thus, there is not only one single muscle working, but e.g. the three heads of the respective muscle. The cooperation of those three oscillating actuators are still not uncovered completely. However, as shown in terms of isometric muscle activity, collaborating muscles and tendons can be synchronized by the neuromuscular system [3,28]. Thereby, short phases without synchronization are alternating with long phases of significant coherence. It is assumed that the three muscle heads should also be able to develop coherent behavior, which is supposed to be controlled by supraspinal motor areas. Since all of those muscle heads insert into one tendon, a superpositioning effect of the tendon is assumed for the Achilles or triceps brachii tendon. Measures of tendons, therefore, could reveal further insights into the quality of motor controlling processes and, in general, into motor control.

The Achilles tendon, e.g., is alternated tightened and released during walking. Due to the impacts of floor during heel strike and the contraction of the triceps surae muscle during push off, the sinew is tightened. The behavior of the tendinous string during and after this impact is influenced by the mentioned active drives of the muscles but also by its passive mechanical properties. The tension and length influences the resonance frequency like it is the case for a chord of a guitar. Thereby, the tendon function as band pass. Therefore, certain frequencies are suppressed and the surrounding soft tissues will have a vibration damping effect. It is therefore assumed that the oscillations of tendons not simply reflect muscular vibrations, but the behavior is highly influenced by the tension and the vibrations of their driving muscles. It is supposed that if the motor control is restricted, changes in the mechanical oscillating behavior of tendons might reflect them and, therefore, investigating those mechanical tendinous oscillations might provide a more functional insight into the properties of tendons. Hence, the non-invasive and easy applicable method of mechanotendography could be a promising option to be applied in further studies investigating the musculoskeletal-system to enlarge the knowledge of the behavior of those relevant bodily structures in healthy and diseased persons and to examine this promising tool for probable applications in diagnostics.

## Conclusion

The repetition trials showed that the used piezo-based measurement system is suited to measure mechanical oscillations reproducibly. It is concluded that the MTG is a reliable and valid tool to measure tendinous oscillations. It seems reasonably transferable to muscular oscillations (mechanomyography).

The methods of mechanotendography and mechanomyography open up possibilities to get insights into the tendinous and motor output, which might reflect the functionality of the neuromuscular system and control. Therefore, the application of this innovative, non-invasive, easy applicable method in further studies dealing with the neuromuscular system is suggested as one practicable approach. It might lead to additional knowledge of pathomechanism and might help in diagnosing impairments of the neuromuscular system and motor control. Further studies should focus and the connection between oscillatory pattern and tensile structure.

## COPYRIGHT REMINDER

All figures, including the pictures, were made in the Neuromechanics laboratory of the University of Potsdam. Therefore, no copyright permission has to be obtained from other sources.

## ACKNOWLEDGMENT

Not applicable.

## Authors contributions

LVS and FNB designed the study. LVS performed the measurements, data processing, analysis and drafted the manuscript. Both authors reviewed the manuscript critically.

## FUNDING

No funding applicable.

## NOMENCLATURE

CA: Cronbach’s alpha
CV: Coefficient of variation
ECG: Electrocardiography
EMG: Electromyography
ICC: Intra class correlation coefficient
M: Mean
MCC: Mean spearman correlation coefficient
MD: Mean distances
MMG: Mechanomyography
MTG: Mechanotendography
MVC: Maximal voluntary contraction
NI: National Instruments
SD: Standard deviation
SNR: Signal to noise ratio

